# Comparative *in vitro* transcriptomic analyses of COVID-19 candidate therapy hydroxychloroquine suggest limited immunomodulatory evidence of SARS-CoV-2 host response genes

**DOI:** 10.1101/2020.04.13.039263

**Authors:** Michael J. Corley, Christopher Sugai, Michael Schotsaert, Robert E. Schwartz, Lishomwa C. Ndhlovu

**Author notes:** Corresponding Author* **Correspondence:** Lishomwa C. Ndhlovu MD, PhD, Division of Infectious Diseases, Department of Medicine, Weill Cornell Medical College, 413E 69^th^ St, New York, NY, USA.

## Abstract

Hydroxychloroquine (HCQ) has emerged as a potential and controversial antiviral candidate therapy for COVID-19. While many clinical trials are underway to test the efficacy of HCQ as a treatment for COVID-19, underlying mechanisms of HCQ in the setting of COVID-19 remain unclear. Hence, we examined differential gene expression signatures of HCQ exposure, *in vitro* SARS-CoV-2 infection, and host signatures of COVID-19 in blood, bronchoalveolar lavage, and postmortem lung to evaluate whether HCQ transcriptome signatures associate with restoration of SARS-CoV-2-related host transcriptional responses. Here, we show that 24 hours of *in vitro* treatment of peripheral blood mononuclear cells(PBMC) with HCQ significantly impacted transcription of 16 genes involved in immune regulation and lipid metabolism. Using transcriptome data from *in vitro* SARS-CoV-2 infected NHBE and A549 cells and PBMC derived from confirmed COVID-19 infected patients, we determined that only 0.24% of the COVID-19 PBMC differentially expressed gene set and 0.39% of the *in vitro* SARS-CoV-2 cells differentially expressed gene set overlapped with HCQ-related differentially expressed genes. Moreover, we observed that HCQ treatment significantly impacted transcription of 159 genes in human primary monocyte-derived macrophages involved in cholesterol biosynthetic process and chemokine activity. Notably, when we compared the macrophage HCQ-related gene lists with genes transcriptionally altered during SARS-CoV-2 infection and in bronchoalveolar lavage of COVID-19+ patients, the *CXCL6* gene was impacted in all three transcriptional signatures revealing evidence in favor of chemokine modulation. HCQ-related transcriptional changes minimally overlapped with host genes altered in postmortem lung biopsies from COVID-19 participants. These results may provide insight into the immunomodulation mechanisms of HCQ treatment in the setting of COVID-19 and suggest HCQ is not a panacea to SARS-CoV-2 infection.

## INTRODUCTION

An outbreak of a novel respiratory disease appeared in Wuhan, China in December 2019[1] capable of human infection and stealth transmission [2,3]. Rapid scientific investigation identified the causative pathogen as a coronavirus named SARS-CoV-2 with similarity to SARS. Clinical observations have shown that the disease arising from SARS-CoV-2 infection termed COVID-19 is characterized by a myriad of symptoms including cough, fever, lethargy, and difficulty breathing[3]. Currently, there remains no effective therapy for COVID-19; hence, a variety of clinical trials are under way to identify an effective COVID-19 therapy. Hydroxychloroquine (HCQ) has emerged as a controversial and potential candidate therapy for COVID-19[4,5]; however, the mechanisms of HCQ antiviral action are unclear. Hence, understanding host biological mechanisms that change during HCQ treatment will be key to understanding HCQ in the setting of COVID-19.

While the efficacy of HCQ as a treatment for COVID-19 remains to be determined, we hypothesized that comparative analyses of HCQ transcriptome signatures and SARS-CoV-2-related host transcriptional responses may provide insight into mechanisms of HCQ treatment. To start to test this hypothesis, we investigated transcriptional signatures of HCQ treated human primary PBMC, human monocyte-derived macrophages, and human plasmacytoid dendritic cells in relation to *in vitro* SARS-CoV-2 infection and COVID-19 participant PBMC, bronchoalveolar lavage, and postmortem lung tissue biopsy transcriptional signatures. These comparative transcriptome findings do not address the question of the efficacy of HCQ in patients with COVID-19, but begin to explore potential mechanisms of HCQ in the setting of SARS-CoV-2 infection.

## RESULTS AND DISCUSSION

### Treatment of resting peripheral blood mononuclear cells with HCQ for 24 hours minimally impacts transcriptome

Due to the immunomodulation effects reported HCQ[6] we surveyed whether *in vitro* treatment of peripheral blood mononuclear cells from healthy people with hydroxychloroquine (10 μM) impacted immune cell gene transcription. We used transcriptome profiling data from three heathy participants PBMC cells that had been either cultured *ex vivo* for 24 hours or treated with hydroxychloroquine for 24 hours (**Supplemental Table 1**). We examined differential gene expression from gene counts using edgeR[7] and corrected for multiple comparisons using the FDR method and observed 16 genes that significantly differed between 24 hour untreated and 24 hour hydroxychloroquine conditions at an adjusted P value of P < 0.05 (**Fig.1a**). The most significant difference was observed for the *IL1R2* gene linked to immune regulation[8], which was downregulated with HCQ treatment. Transcription of *IL1R2* has been reported to be significantly upregulated in monocytes from individuals with chronic graft-versus host disease[9] suggesting HCQ may exert anti-inflammatory effects. Notably, we observed that the chemokine *CCL22* gene was significantly down regulated after 24 hours of HCQ treatment as compared to the untreated condition (**Fig.1a**). *CCL22* codes for a CC cytokine that is involved in chemotactic activity for monocytes, dendritic cells, natural killer cells, and activated T cells. Other genes that were significantly differently expressed comparing the untreated and HCQ treated conditions included *SEMA3G*, *FN1*, *MSMO1*, *ARHGEF28*, *SLC1A2*, *PHOSPHO1*, *ENC1*, *CPNE6*, *SCD*, *NDRG2*, *LDLR*, *A2M*, *AQP9*, and *CYP51A1*. Pathway analyses of this gene set using the Enrichr gene enrichment analysis tool[10] showed that the top pathways involved lipid and lipoprotein metabolism, SREBF and miR-33 in cholesterol and lipid homeostasis, lipoprotein metabolism, and steroid biosynthesis (**Supplemental Data 1**). These *in vitro* data suggest that 24 hour *in vitro* treatment with HCQ minimally impacts immune gene transcription and support findings that HCQ has potential immunomodulatory and lipid metabolism effects[11]. Specifically, previous work from *in vivo* studies has reported that HCQ treatment significantly decreased serum TNF-A and increased IL-10 levels in women following treatment with 400mg/per day orally[12] and inhibited IL-1β production from stimulated human neutrophils[13]. In the context of COVID-19 infection, clinical reports indicate severe patients exhibit a well-known immune cytokine storm that leads to multi-organ failure[14,15]. Ongoing clinical trials will see if in addition to hypothesized *in vivo* viral impacts of HCQ whether this drug also averts the likelihood for a cytokine storm.

### HCQ-related transcriptional changes in PBMC minimally overlap with host genes altered during *in vitro* SARS-CoV-2-infection

While there remains a lack of treatments for COVID-19, HCQ has received increased attention due to data from *in vitro* experiments suggesting antiviral activity[4]. In an effort to examine whether the HCQ-related transcriptional changes in PBMC cells related to *in vitro* SARS-CoV-2 impacted gene transcription upon infection, we examined the overlap of the 16 differentially expressed HCQ genes with recent transcriptional data from *in vitro* SARS-CoV-2 infection of NHBE cells and A549 cells (**Supplemental Table 1**). This comparative analysis identified only two overlapping genes (**Fig.1b**), NDRG Family Member 2 (*NDRG2)* and Low Density Lipoprotein Receptor (*LDLR)* genes, which were transcriptionally altered in the HCQ treatment of PBMC transcriptional dataset (**Fig.1c,f**) and in the SARS-CoV-2 *in vitro* infection of NHBE cells transcriptional dataset (**Fig.1d,g**). Interestingly, the *NDRG2* and *LDLR* genes were not altered upon SARS-CoV-2 *in vitro* infection of A549 cells (**Fig.1e,h**), suggesting different *in vitro* SARS-CoV-2 infection models vary in transcriptional responses which will be critical for identifying relevant host genes impacted by viral infection. The *NDRG2* gene has been identified as a novel factor for the regulation of IL-10 in myeloid cells[16] and a factor in macrophage polarization[17], suggesting HCQ may impact myeloid cell inflammation that has been hypothesized as a driver of severe COVID-19[18]. We observed that *NDRG2* was significantly upregulated in the SARS-CoV-2 *in vitro* infection dataset (**Fig.1d**) and significantly downregulated in the *in vitro* hydroxychloroquine treatment dataset (**Fig.1c)**, suggesting that HCQ may restore host transcription of *NDRG2* and exert anti-inflammatory effects. We found that the *LDLR* gene was significantly upregulated in both the *in vitro* hydroxychloroquine treatment and SARS-CoV-2 infection datasets (**Fig.1f,g**), suggesting both HCQ treatment and SARS-CoV-2 infection alter lipid metabolism transcription. We acknowledge these observations are associative and further casual studies are needed. Previous work has shown HCQ impacts lipid profiles[19] by potentially altering intracellular pH leading to lipid uptake and increased receptor expression. Whether HCQ exerts immunomodulator effects by altering transcription of cell lipid metabolic pathways needs further study.

### HCQ-related transcriptional changes in PBMC minimally overlap with host genes altered during human SARS-CoV-2-infection

Next, we sought to perform a more direct comparison of transcriptional differences observed by HCQ treatment of primary PBMC cells with transcriptional differences in a dataset of PBMC samples from SARS-CoV-2 infected individuals compared to uninfected controls. We analyzed RNA-seq data of PBMC samples from three male individuals infected with SARS-CoV-2 and three control individuals without SARS-CoV-2 infection obtained from China (**Supplemental Table 1**)[20]. We first sought to confirm the observed immunological changes observed during COVID-19 disease in peripheral blood by others[15,21–24]. We utilized the bulk transcriptome PBMC data and estimated the abundances of 22 immune cell types using the CIBERSORT analytical deconvolution tool[25]. Similar to other reports, we observed a drastic expansion of monocyte cells in peripheral blood of COVID-19 participants (75.3%, 46.4%, and 57.4%) compared to controls (19.8%, 5.6%, and 26.8%) using an estimation of member cell types in PBMC based on gene expression data (**Fig. 2a, Supplemental Figure 2**). Also, we saw that CD4 memory resting cells and NK cells resting drastically reduced in all three COVID-19 participants compared to controls (**Fig. 2a, Supplemental Figure 2**). The estimates of all 22 immune cell types in PBMC from COVID-19 and controls are presented in **Supplemental Figure 2**. Next, we examined the overlap of HCQ-related transcriptional changes we had observed in PBMC cells with the differentially expressed gene set related to COVID-19 in PBMC. Of the 2,082 differentially expressed genes related to COVID-19, we identified that only 5 genes overlapped between the gene set of HCQ-related transcriptional changes and COVID-19-related transcriptional changes in PBMC (**Fig. 2b**). This accounted for only 0.24% of the host related COVID-19 transcriptional changes in PBMC, suggesting HCQ minimally restores/impacts COVID-19-related transcriptional immune changes. In the HCQ PBMC transcriptional dataset, we observed that the *IL1R2, FN1,* and *A2M* were significantly downregulated and *LDLR* and *SCD* gene were significantly upregulated (**Fig.2c**). In the COVID-19 PBMC transcriptional dataset, we observed that the *LDLR* gene was significantly upregulated similar to the SARS-CoV-2 *in vitro* infection dataset for NHBE (**Fig.2d**). Additionally, in the COVID-19 PBMC dataset, the *IL1R2*, *FN1*, and *SCD* genes were significantly upregulated compared to a significant downregulation of the *A2M* gene (**Fig.2d**). These results should be taken with caution as studies are needed comparing PBMC cells from COVID-19 participants before and after HCQ treatment. Additionally, whether the directionally of transcriptional change in PBMC cells due to HCQ treatment or related to COVID-19 has any immunomodulatory effect leading to clinical benefits requires further study. This comparative analysis is limited and does not address whether HCQ has antiviral effects by reducing SARS-CoV-2 virus entry through impacting cell lipid metabolism pathways since PBMC cells are not a main target for SARS-CoV-2 infection and viral replication. We preliminarily confirmed that PBMC cells do not harbor SARS-CoV-2 virus by mapping RNA-Seq reads from COVID-19 PBMC samples and identifying no significant reads aligning to the SARS-CoV-2 reference (**Supplemental Figure 3b**). This was also observed by Xiong et al. [20]. In contrast, we observed viral mapping of reads in the Blanco-Melo et al. *in vitro* SARS-CoV-2 infection dataset of A549 and NHBE cells (**Supplemental Figure 3c,d)**. Notably, the *in vitro* SARS-CoV-2 infection dataset contains a 3’ bias of read alignment compared to the COVID-19 PBMC dataset. Recent *in vitro* work suggest SARS-CoV-2 infects T lymphocytes[26]. Whether this is observable in COVID-19 patient remains to be determined. We assume HCQ treatment of A549 or NHBE cells during SARS-CoV-2 infection would dramatically reduce the viral mapping detected by RNA-Seq. Our observations of overlapping transcriptional changes provides candidate genes to further study for addressing host immune-related mechanisms of action of HCQ in the setting of COVID-19.

### Treatment of human primary monocyte-derived macrophages with HCQ impacts chemokine and lipid metabolism-related genes

Previous research in SARS suggested that SARS-CoV regulates immune function-related gene expression in myeloid cells[27] and a report in a SARS-CoV-infected mouse model showed that dysregulated inflammatory and interferon responses of monocyte/macrophage cells caused lethal pneumonia[28]. Clinical data about COVID-19 suggest that peripheral innate immune monocyte cells and lung tissues macrophage cells are critically involved in host response and severe disease course[29]. Together, these findings suggest that modulation of myeloid cells may have an impact on SARS-CoV-2-related host responses. Interestingly, research on HCQ suggest that it may impact TLR signaling, antigen presentation, and alter macrophage function[30]. HCQ is also proposed to indirectly have anti-inflammatory effects upon the immune system. Hence, we examined transcriptional differences in human primary CD14+ monocyte MCSF-differentiated macrophage conditions treated with HCQ for 24 hours (**Supplemental Table 1**). Differential gene expression analyses showed that 159 genes were differentially expressed comparing the untreated and HCQ treated conditions corrected for multiple comparisons using the FDR method at P < 0.05 (**Supplemental Data 1**). Among these 159 genes, gene enrichment analyses showed the top GO biological process was cholesterol biosynthetic process, GO molecular function chemokine activity, and GO cellular component integral component of plasma membrane (**Fig.3a-c, Supplemental Data 1**). Specifically, HCQ altered expression of *LDLR*, *CXCL6*, *CCL18*, *CCL5*, and *CXCL3* genes (**Fig.3d-h**). To extend our findings of HCQ impacting immune chemokine activity in innate immune cells, we examined transcriptomes of plasmacytoid dendritic cells (pDC) from four healthy individuals that had been either stimulated with RNA-IC for 6 hours only or stimulated with RNA-IC for 6 hours in the presence of HCQ (**Supplementary Table 1**). We observed that *CXCL10*, *CXCL11*, *CCL4*, and *CXCL9* genes were significantly downregulated in stimulated HCQ treatment compared to the only stimulated condition (**Supplemental Figure 4**). Together, our findings in macrophages and pDC cells suggest HCQ impacts certain cell type specific chemokine signaling pathways and likely modulates immune inflammation.

### HCQ-related transcriptional changes in macrophages overlap with 7 host genes altered during *in vitro* SARS-CoV-2-infection and in bronchoalveolar lavage from COVID-19 participants

In our comparative analyses for the macrophage HCQ dataset, we first examined whether the HCQ-related transcriptional changes in macrophage cells related to SARS-CoV-2 impacted gene transcription from *in vitro* SARS-CoV-2 infection of NHBE cells and A549 cells (**Supplemental Table 1**). This comparative analysis identified 15 overlapping genes including *LDLR*, *CXCL6*, *GPNMB*, *HMGCS1*, *IDI1*, *C1S*, *FDPS*, *MMP1*, *PLAU*, *MAFF*, *ITGB3*, *BHLHE40*, *TNC*, *CXCL3*, *SLC7A8* (**Fig.4a**). Next, we sought to compare the HCQ-related transcriptional changes in human primary macrophage cells with transcriptional data from COVID-19 participants and in a similar cell type. Hence, we analyzed transcriptome data from bronchoalveolar lavage samples obtained from two individuals with COVID-19 and three uninfected individuals (**Supplemental Table 1**). Bronchoalveolar lavage (BAL) samples predominantly contain macrophages with only a minor population of neutrophils, lymphocytes, and eosinophils. We used CIBERSORT to confirm that the majority population in the transcriptome dataset was macrophage cell types (**Supplemental Figure 5**). Additionally, we confirmed that the BAL transcriptome dataset from COVID-19 participants contained high levels of SARS-CoV-2 virus by mapping reads to the viral reference (**Supplemental Figure 3a**). The bronchoalveolar lavage COVID-19 transcriptome dataset contained 5,304 differentially expressed genes comparing COVID-19 to uninfected individuals. Our comparative analysis of the macrophage HCQ-treated gene set with the COVID-19 lavage transcriptional dataset identified 45 overlapping genes accounting for only 0.84% of differentially expressed genes related to COVID-19 (**Fig.4a**). Seven of the 45 genes overlapped with the *in vitro* SARS-CoV-2 infection transcriptional dataset and included *CXCL6*, *IDI1*, *MMP1*, *PLAU*, *MAFF*, *BHLHE40*, and *SLC7A8* genes (**Fig.4a-d)**. Our data suggest that studies examining the regulation of these genes and relationship to SARS-CoV-2 infection and host immune responses are warranted.

### HCQ-related transcriptional changes in macrophages minimally overlap with 4 host genes altered in postmortem lung biopsies from COVID-19 participants

Immune regulation and defenses against viral pathogens occur within in the lung. We sought to examine whether the differential gene expression signatures of HCQ in immune cells associated with a COVID-19-related transcriptional profile in the lung. We analyzed a limited transcriptomic data from postmortem lung biopsies obtained from two male participants with COVID-19 over the age of 60 years and two uninfected age-matched control male participants (**Supplemental Table 1**). The COVID-19 and uninfected control participants were confirmed to not have been on HCQ treatment. As expected, we also detected 108 viral SARS-CoV-2 reads in the bulk RNA-Seq lung dataset of one of the COVID-19 individuals and 7 viral SARS-CoV-2 reads in the second COVID-19 participant (**Supplemental Figure 6)**. These results are limited by the low sequencing depth and 3’ bias but do offer an interesting means to confirm COVID-19 infection by RNA-seq screening in postmortem samples. Our comparative analysis of the both PBMC and macrophage HCQ-treated gene sets with the COVID-19 postmortem lung transcriptional dataset identified only 5 overlapping genes (**Fig.5a**). This overlap included *CCL18*, *FOS*, *MMP1*, *ITGB3*, and *AQP9* genes. Notably, our gene ontology analysis of the COVID-19 postmortem lung dataset showed the top GO Biological Process as negative regulation of viral genome replication, GO Molecular Function as chemokine activity, and GO Cellular Component as secretory granule lumen. Additionally, we used CIBERSORT[25] to examine the immune cell populations in COVID-19 lung compared to uninfected lung (**Supplemental Figure 7**). This deconvolution analysis showed some interesting initial differences of interest for further study. First, the relative fraction of CD4 memory resting T cells was 15% and 19% in the uninfected lungs compared to only 3% and 4% for the COVID-19 lung. Second, the relative fraction of neutrophils was 23% and 13% in COVID-19 lung compared to 0% and 8% for the uninfected lung. These results support the notion of continual neutrophil influx into the lung in the setting of COVID-19 during sustained and persistent inflammatory cytokine and chemokine secretion[31]. Lastly, in support of the possibility of mast cell activation in severe cases of SARS-CoV-2 infection, we observed that the relative fraction of mast cells activated was 59% and 45% in COVID-19 lung compared to 37% and 0% for uninfected. While these results are limited by the low sample size, they do provide evidence for certain areas of future investigation. We suspect that transcriptome data from HCQ-treated COVID-19 individual’s lung is needed to conclude that HCQ does not dramatically restore SARS-CoV-2 induced lung transcriptome alterations.

In this study, we observed that *in vitro* HCQ treatment of immune cells alters gene transcription which is likely cell type-specific and context dependent. Notably, these alterations were more apparent in macrophage cells compared with bulk PBMC cells treated with HCQ. Our study is a start to address mechanisms of action for whether HCQ is an effective therapy for COVID-19. While these findings are of interest to studies of HCQ and COVID-19, transcriptional studies need to be conducted on *in vivo* treatment of COVID-19 participants before and after HCQ treatment. The mechanisms of action of HCQ in the setting of viral infection remains an area still under investigation and a research area containing contrasting findings. This is supported by previous HCQ research in the setting of HIV that reported HCQ treatment drastically reduced immune activation in HIV-infected individuals[32]. In contrast to these findings, a randomized double blind placebo controlled trial of 400 mg of HCQ failed to find any difference in T cell activation and reported increased viral replication[33]. While *in vitro* data of SARS-CoV-2 provide compelling rationale for suggesting clinical efficacy of HCQ[4,34], *in vivo* data and randomized double blinded placebo COVID-19 studies of HCQ are needed to account of context-dependent effects of the host being administered HCQ. Further work will need to determine whether the mechanism of action of HCQ is directly impacting SARS-CoV-2 or indirectly impacting the immune system and hence indiscriminately improving COVID-19 clinical outcomes. Unfortunately, at this moment HCQ is not a panacea for COVID-19 and alternative therapies are urgently needed.

## CONTRIBUTIONS

MJC and LCN equally contributed to the concept and writing of the manuscript. CS contributed to RNA-Seq analyses and SARS-CoV-2 viral detection in bulk RNA-Seq data. MS edited manuscript and contributed to concept. RES contributed postmortem RNA-Seq data for analysis and edited manuscript.

## ETHICS DECLARATIONS

### Ethics approval and consent to participate

Public Data was obtained from de-identified human participants for this IRB exempt study.

### Consent for publication

Not applicable.

## ACKNOWLEDGEMENTS

We would like to acknowledge the authors that deposited raw data for analysis.

## METHODS

### Comparative analysis strategy

Our comparative analysis utilized Excel, R, and Graphpad Prism software. RNA-Seq gene expression raw FASTQ or gene count data was obtained from the Gene Expression Omnibus (GEO) and BigD database.

### Data

The RNA-Seq gene expression data are listed in Supplemental Table 1. Notes on sample selection are included in the Supplemental Table 1.

### Differential Expression

RNA-Seq count data was read into R using Bioconductor and rowSums >1 were filtered to remove lowly expressed genes. Differential expression was conducted with edgeR and adjusted for P value using the FDR method option. Heatmaps and unsupervised clustering utilized the pheatmap R package. Rows were scaled by calculating the Z score of expression values for each sample. Clustering utilized the complete method.

### Enrichr Gene List Enrichment Analysis

We used the Enrichr integrative web-based analysis tool to perform a gene list enrichment (amp.pharm.mssm.edu/Enrichr) of the annotated differentially expressed gene sets[10].

### RNA-Seq Cell Deconvolution

COVID-19 and uninfected control gene expression data from bulk PBMC, alveolar lavage, and postmortem lung biopsies were analyzed using the CIBERSORT [25]analytical tool to provide an estimate of the abundances of immune cell types in the mixed cell/tissue population. The existing 22 cell type immune signature matrix was used for reference cell types for analysis.

### SARS-CoV-2 Viral Detection in Bulk RNA-Seq Data

Reads were aligned with STAR[35] 2.7.3a in two-pass quantification mode. A custom reference was generated comprising Ensembl GrCh38 release 98 and SARS-CoV-2 2019-nCoV/USA-WA1/2020 (NCBI Nucleotide MN985325.1) to detect viral RNA. Tabulated counts were checked for expected stranded-ness and concatenated in R. Normalization, gene filtering, and differential expression analysis was done edgeR[7]. Genes were filtered for a minimum read count above 10 and log fold change greater than 1. Counts were annotated with AnnotationHub.

---

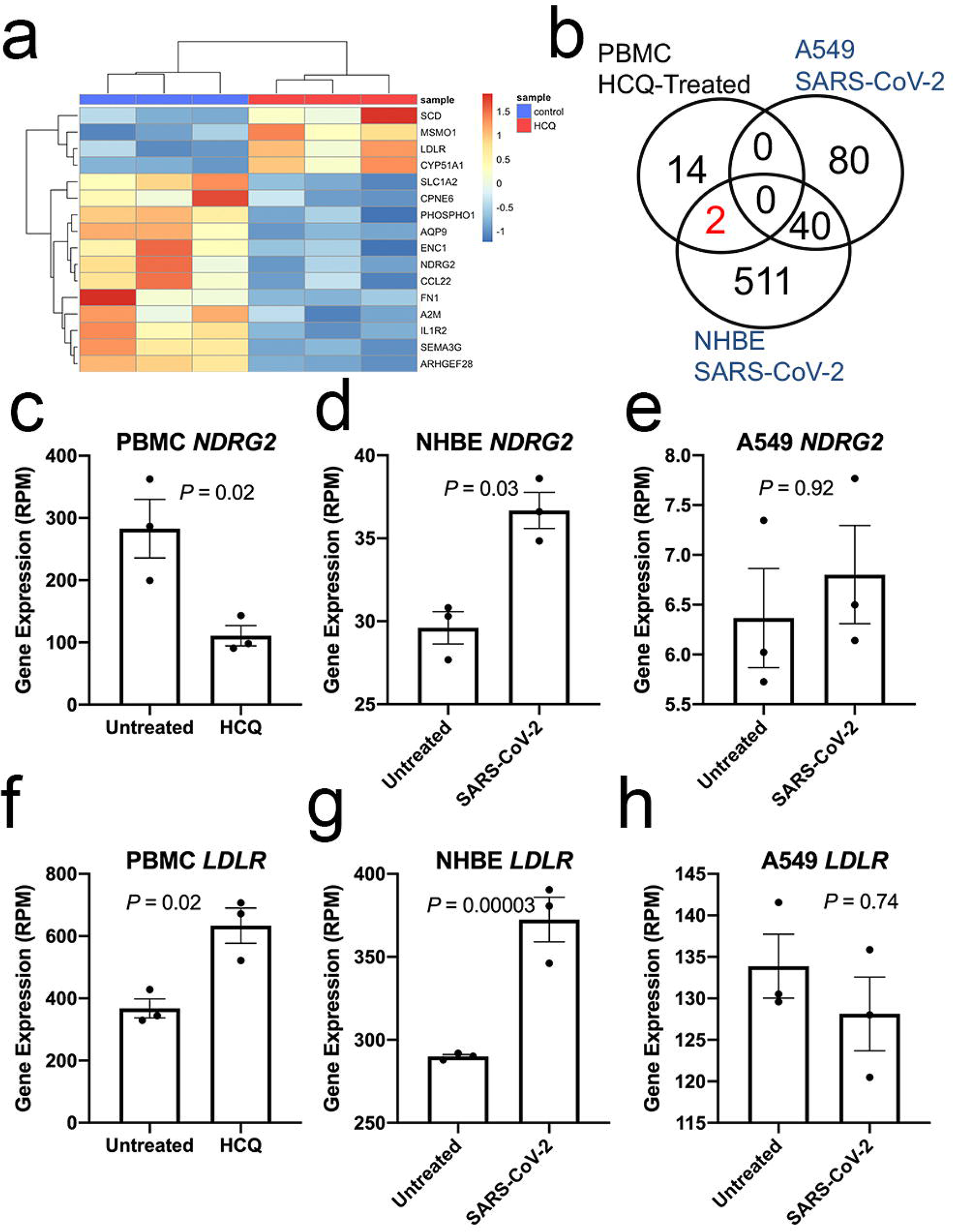

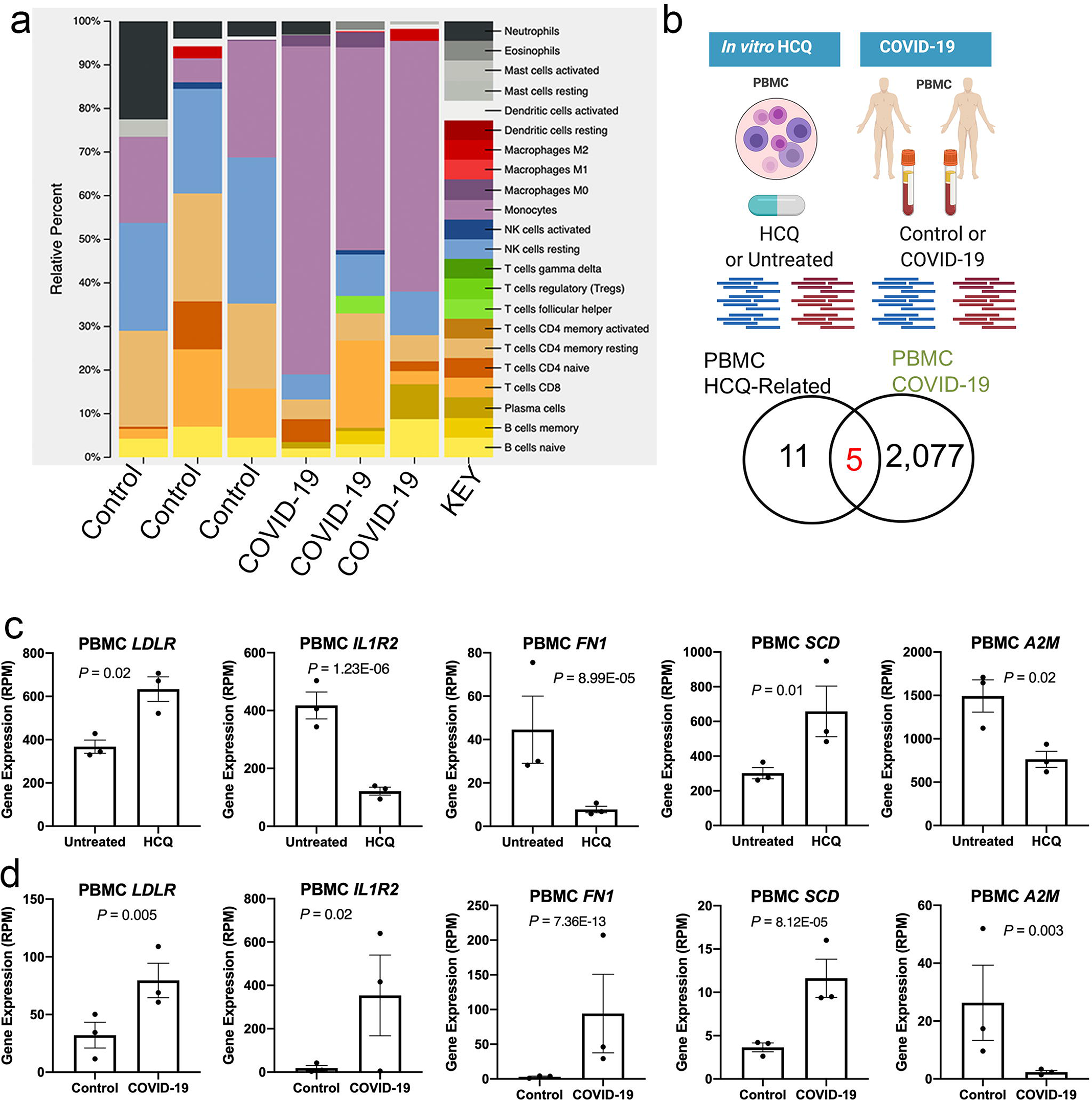

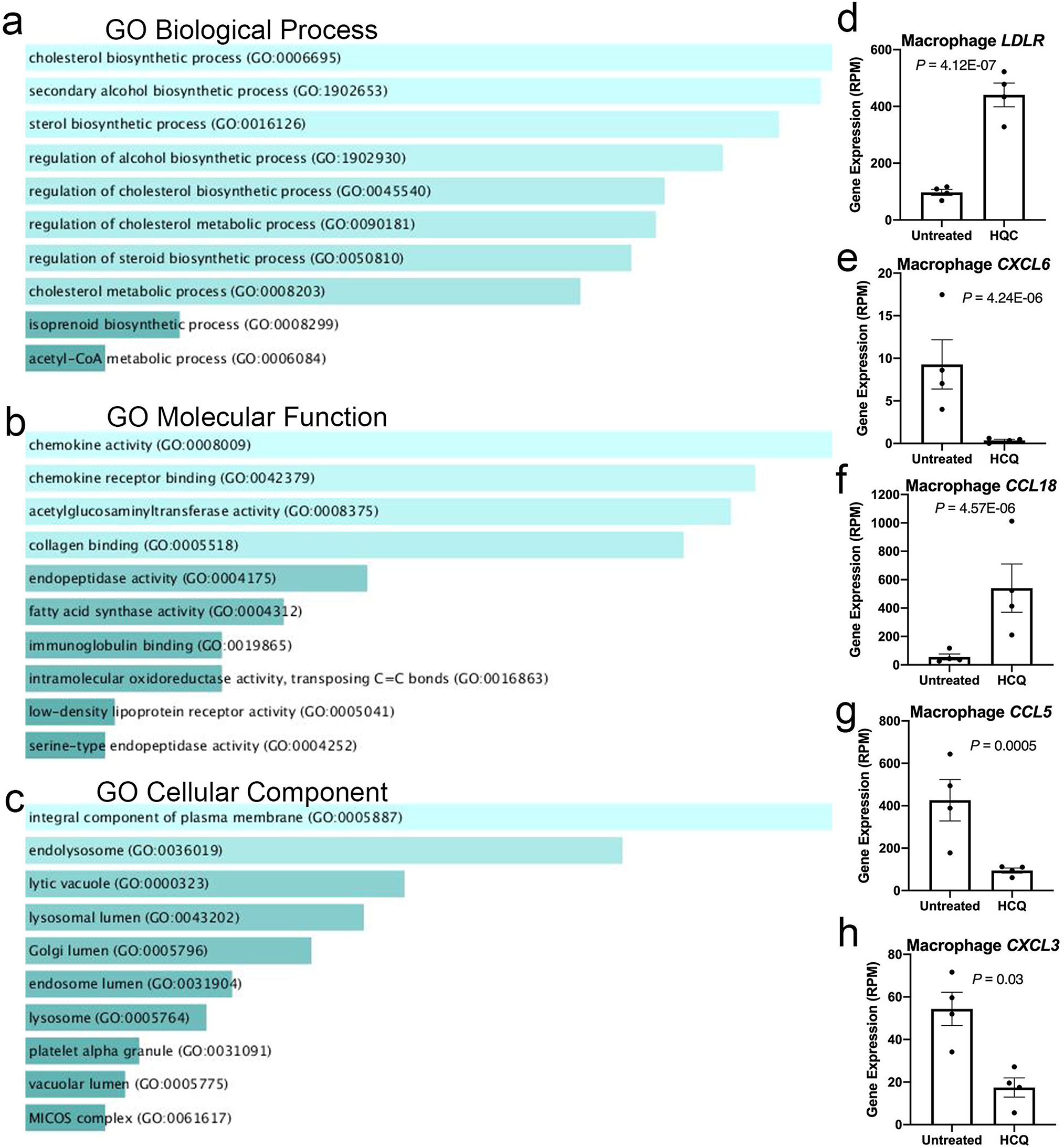

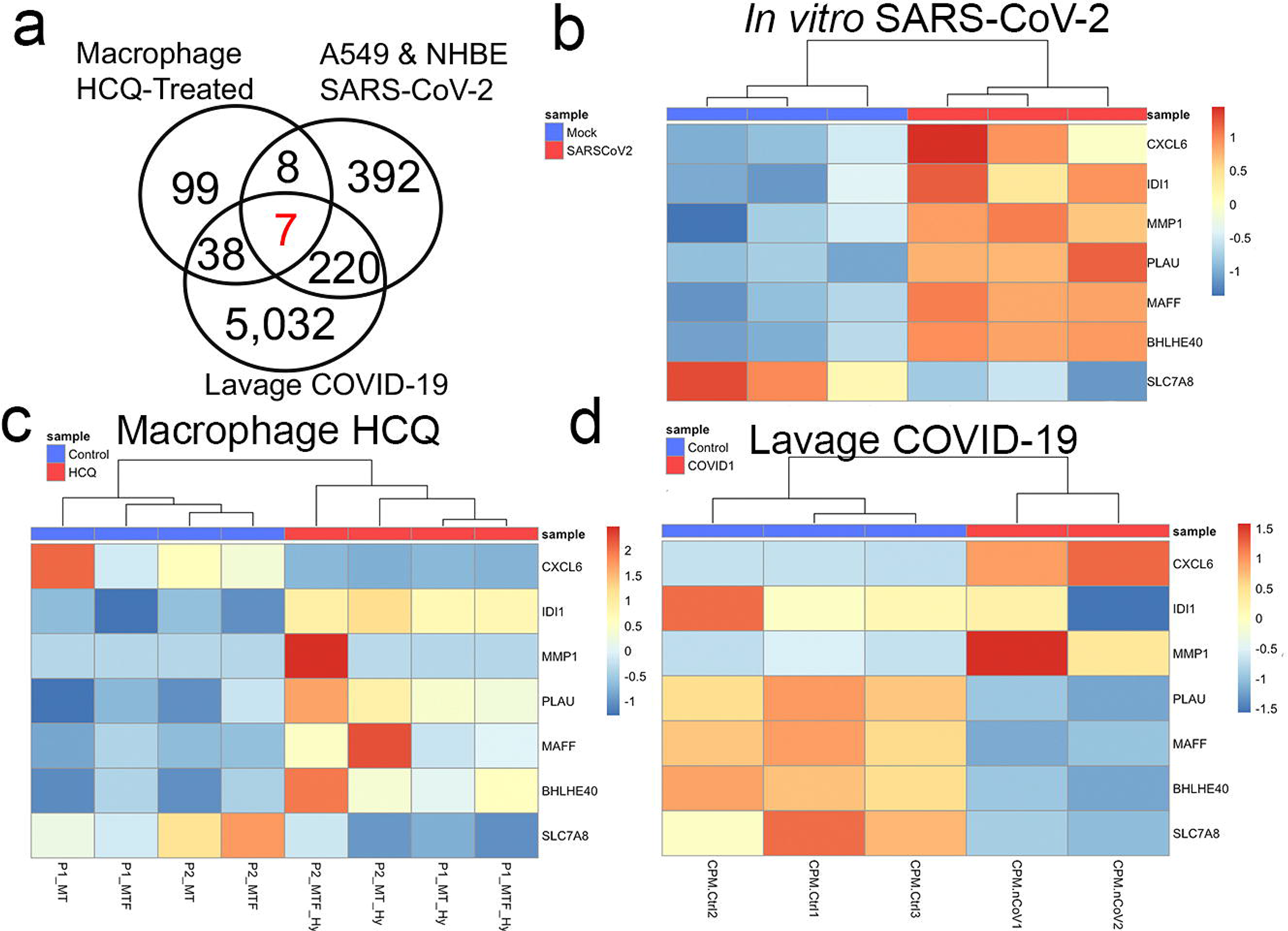

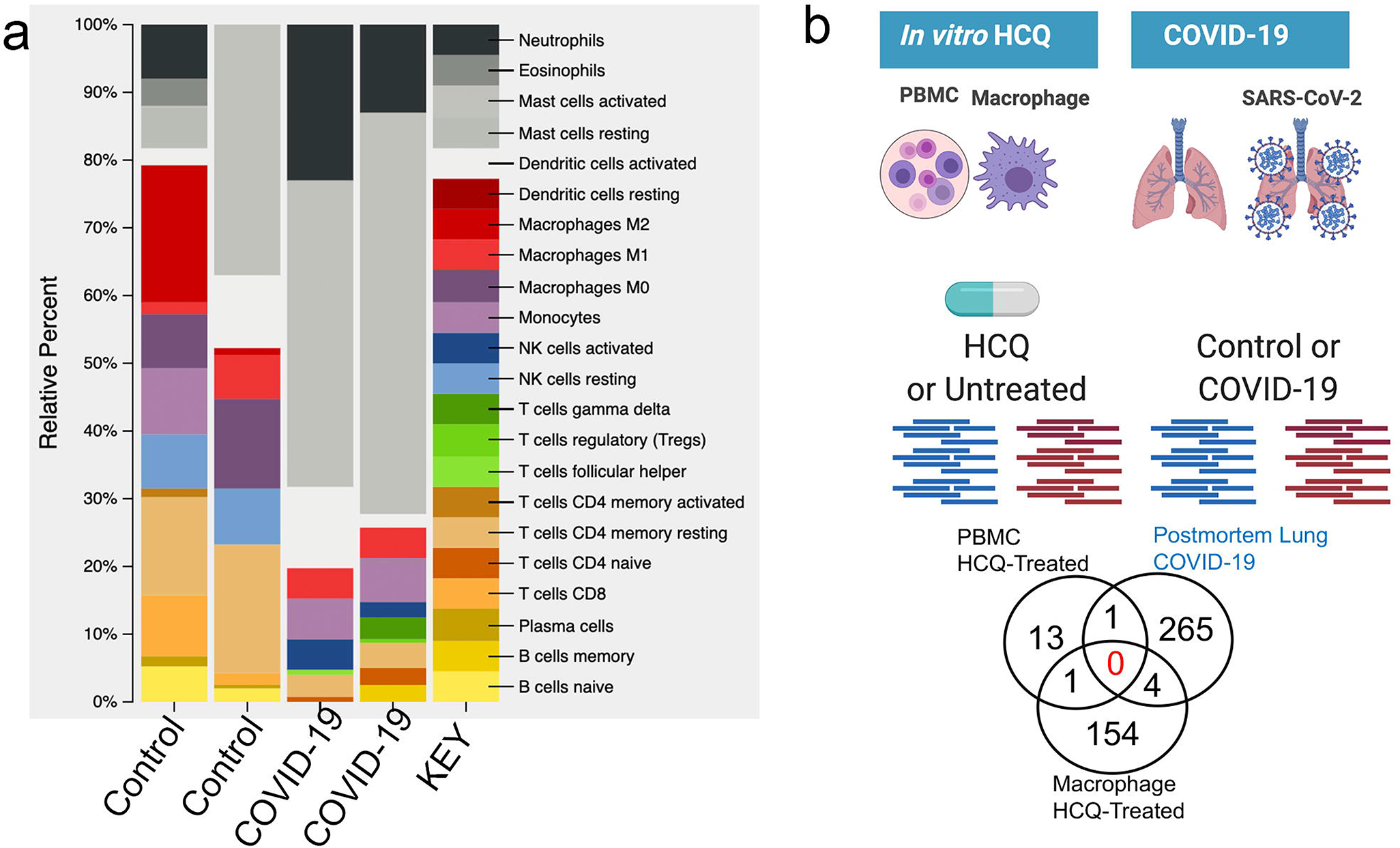

