## supplementalfigurestables for "Comparative *in vitro* transcriptomic analyses of COVID-19 candidate therapy hydroxychloroquine suggest limited immunomodulatory evidence of SARS-CoV-2 host response genes"

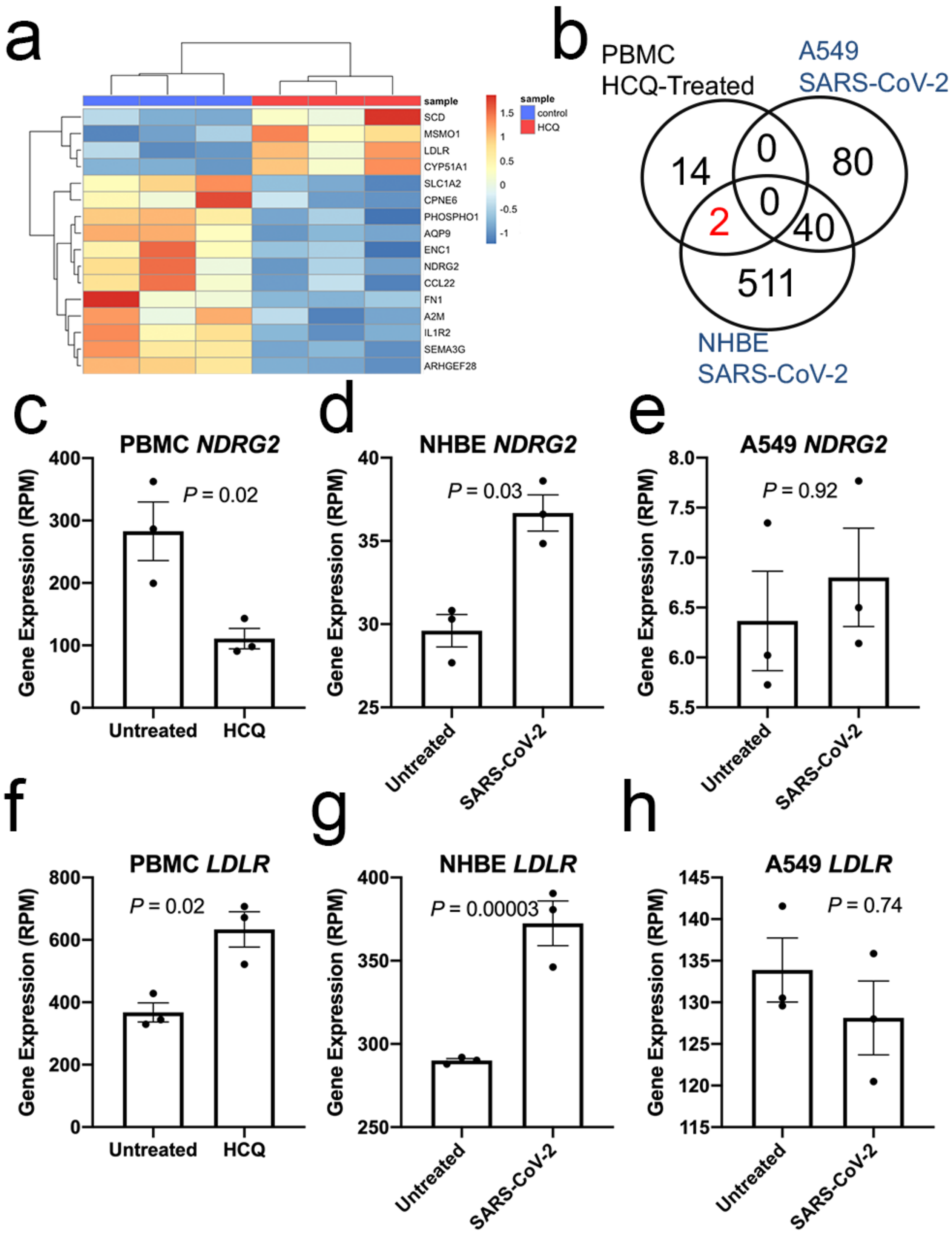


**Figure 1. HCQ-related transcriptional changes minimally overlap with host genes impacted by *in vitro* SARS-CoV-2-infection.** a. Heatmap of differentially expressed genes in untreated (blue) and HCQ treated (red)human PBMC cells. Rows scaled by Z-score. b. Comparison of differentially expressed genes in HCQ treated PBMC (GSE74235), NHBE SARS-CoV-2 infected (GSE147507), and A549 infected cells (GSE147507). Venn diagram showing the number of genes overlapping. c-h. Normalized gene expression in primary PBMC cells either untreated for 24 hours or treated *in vitro* with HCQ (20 μM).

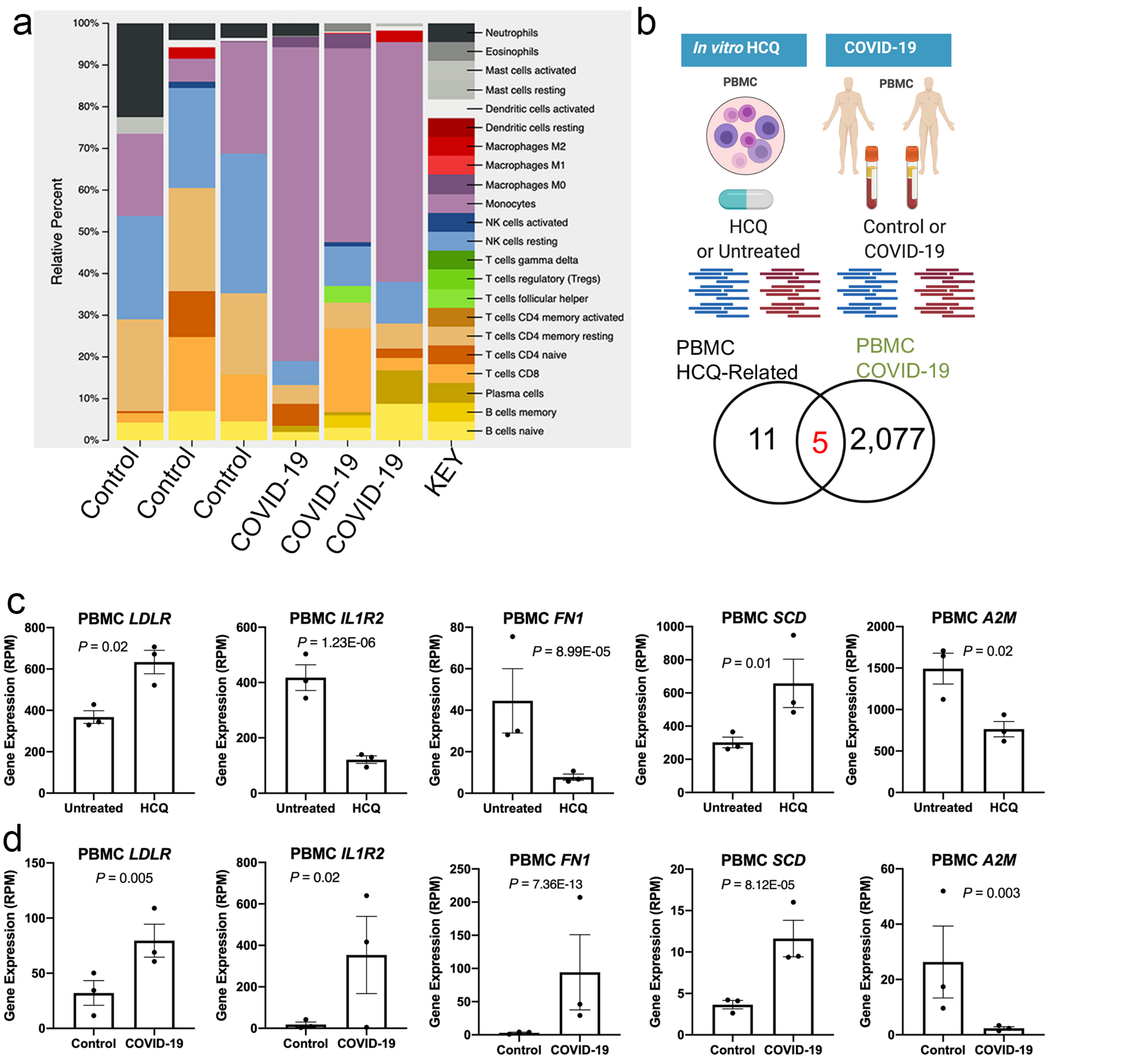


**Figure 2. HCQ-related transcriptional changes minimally overlap with host genes impacted by *in vivo* SARS-CoV-2-infection.** a. Bar graph of relative percent of immune cell types in PBMC samples from uninfected control and SARS-CoV-2 infected (COVID-19) participants deconvoluted from RNA-seq. Color key shown to represent each cell type. b. Comparative analysis of differentially expressed gene signatures of HCQ treatment of PBMC cells and COVID-19. Venn diagram showing overlap of differentially expressed genes. c,d. Normalized gene expression of untreated and HCQ treatment of PBMC cells and uninfected control participants and COVID-19 participants.

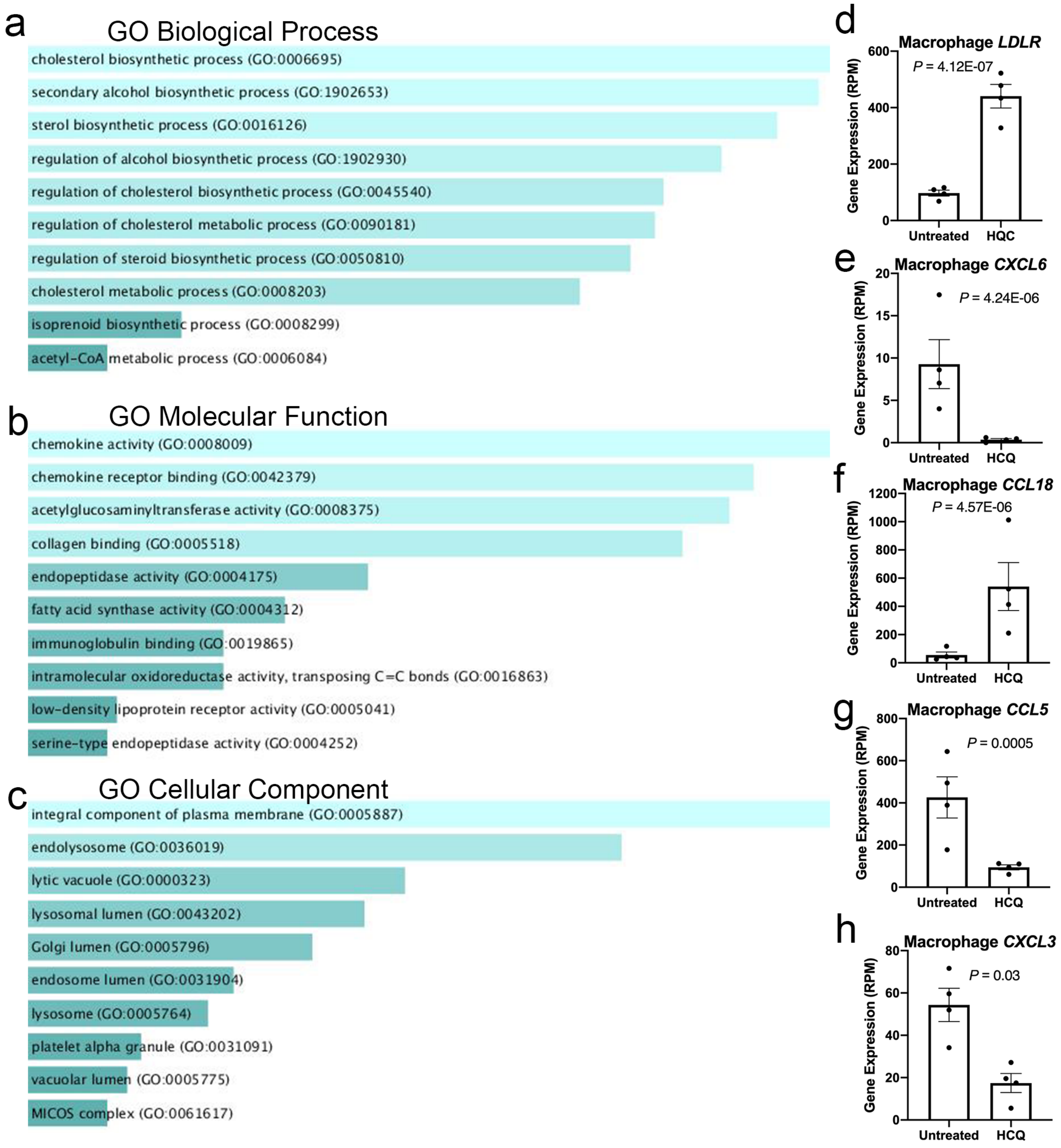


**Figure 3. HCQ-related transcriptional changes in human primary macrophage cells.** a-c. Gene ontology analysis of Biological Process, Molecular Function, and Cellular Component of the 159 genes differentially expressed in HCQ treated macrophage cells. Bar graphs sorted by p-value ranking. d-h. Bar graph of normalized gene expression of *LDLR*, *CXCL6*, *CCL18*, *CCL5*, and *CXCL3* genes differentially expressed in HCQ treatment compared to untreated condition.

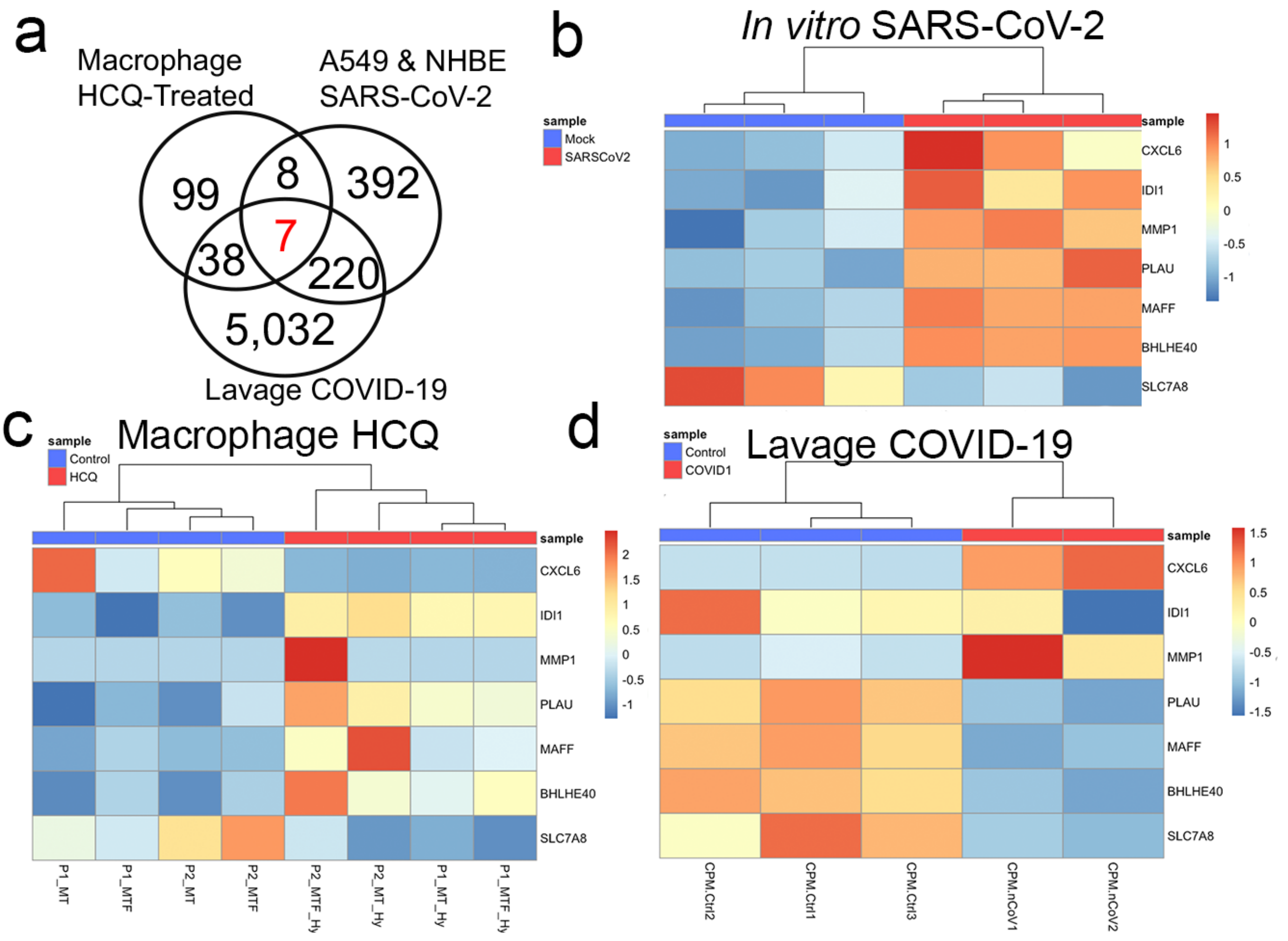


**Figure 4. HCQ-related transcriptional changes in macrophage cells moderately overlap with COVID-19-related transcriptome signature in bronchoalveolar lavage and *in vitro* SARS-COV-2 infection.** a. Comparative analysis of differentially expressed gene signatures of HCQ treatment of macrophage cells and COVID-19-related transcriptome signature in bronchoalveolar lavage and *in vitro* SARS-COV-2 infection. Venn diagram showing overlap of differentially expressed genes. b. Heatmap of overlapping 7 genes from *in vitro* SARS-CoV-2 dataset, c. HCQ treated macrophage dataset, and d. COVID-19 bronchoalveolar lavage dataset. Rows scaled by Z-score

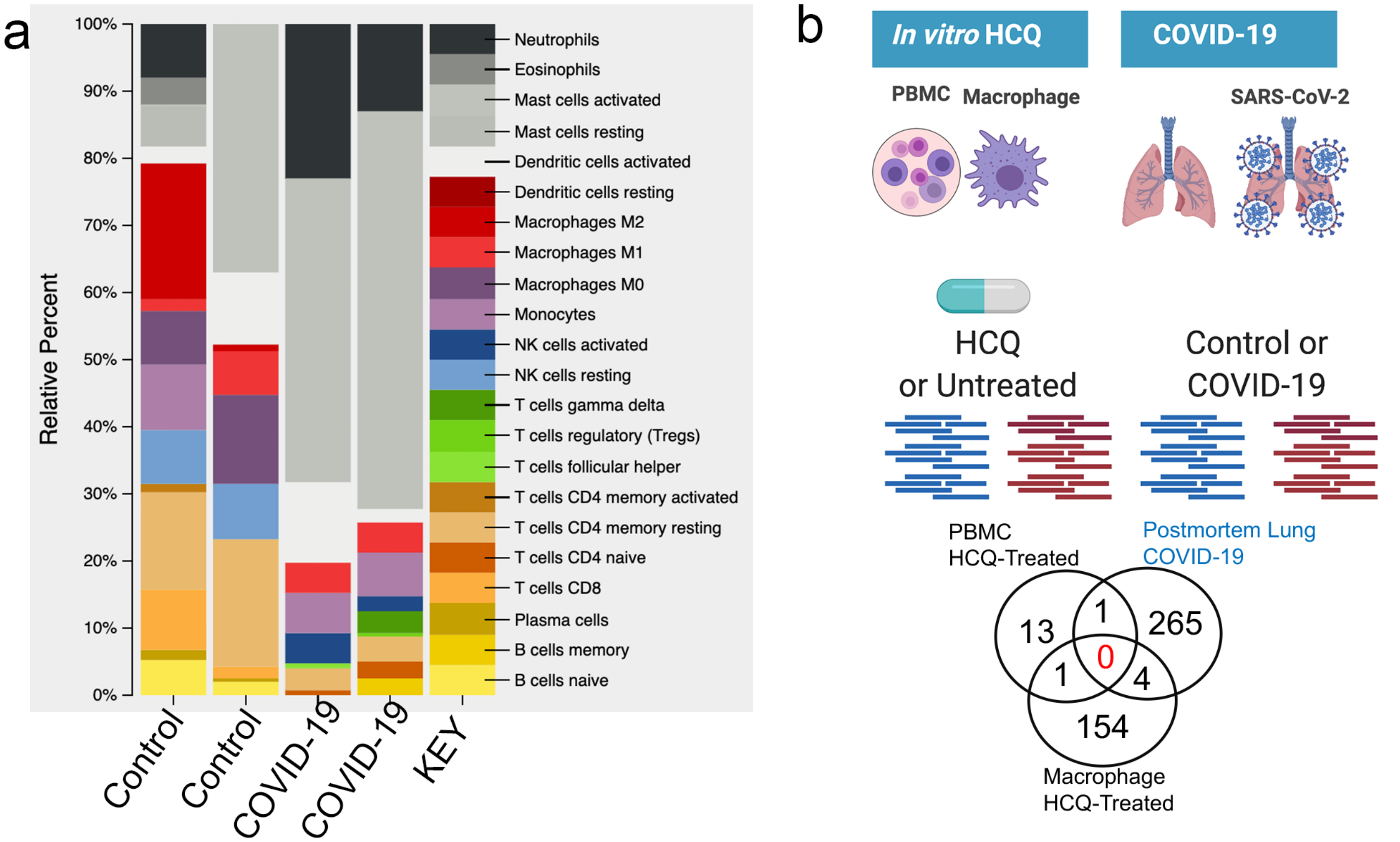


**Figure 5. HCQ-related transcriptional changes minimally overlap with host genes impacted by SARS-CoV-2-infection in postmortem lung.** a. Bar graph of relative percent of immune cell types in postmortem lung samples from uninfected control and SARS-CoV-2 infected (COVID-19) participants deconvoluted from RNA-seq. Color key shown to represent each cell type. b. Comparative analysis of differentially expressed gene signatures of HCQ treatment of PBMC and monocyte-derived macrophage cells and postmortem lung COVID-19. Venn diagram showing overlap of differentially expressed genes.

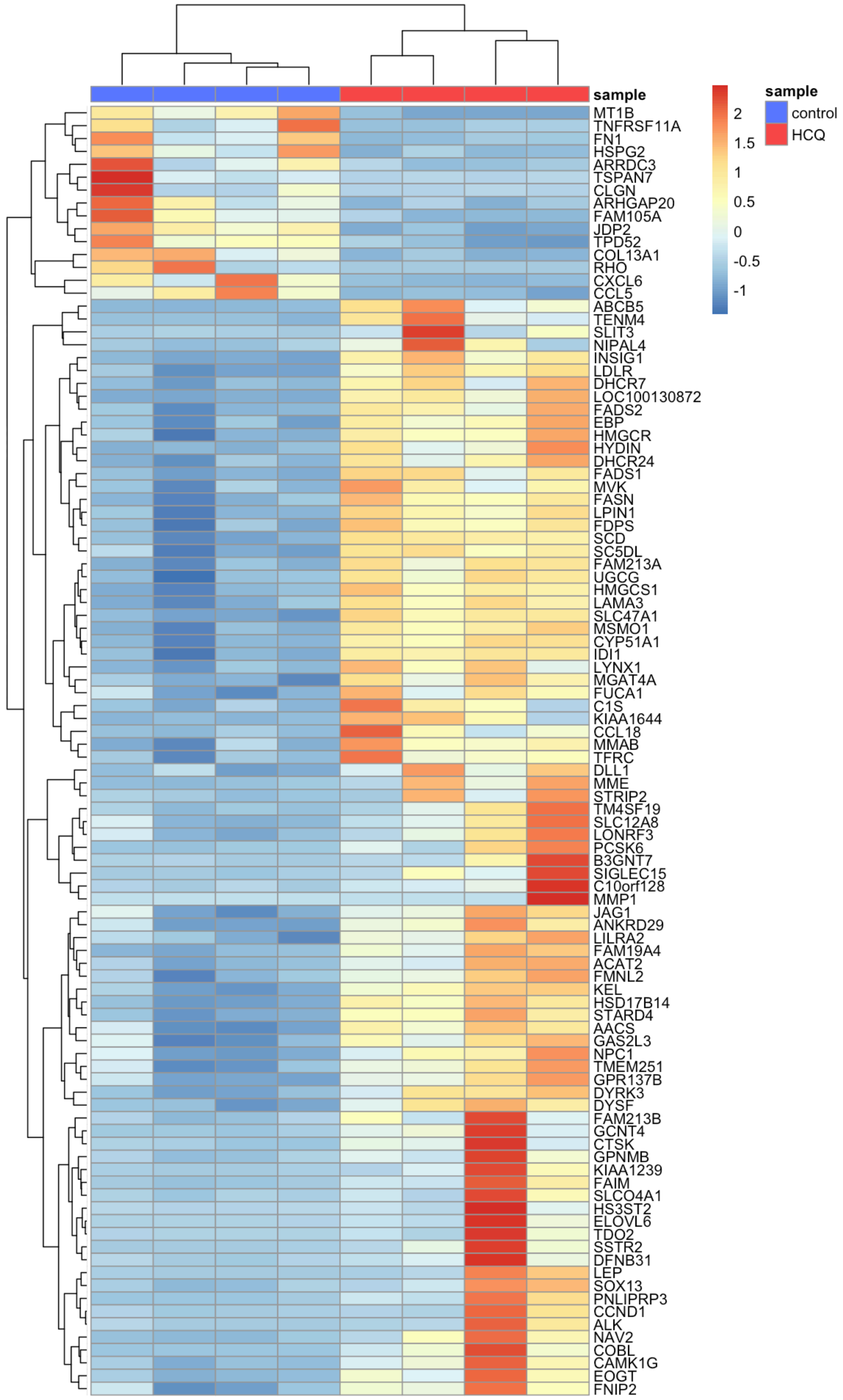


**Supplemental Figure 1. Heatmap of differentially expressed genes in untreated (blue) and HCQ treated (red)human macrophage cells. Rows scaled by Z-score.**

| Input Sample | B cells naive | B cells memory | Plasma cells | T cells CD8 | T cells CD4 naive | T cells CD4 memory resting | T cells CD4 memory activated | T cells follicular helper | T cells regulatory (Tregs) | T cells gamma delta | NK cells resting | NK cells activated |
|---|---|---|---|---|---|---|---|---|---|---|---|---|
| CPM-N1 | 0.043 | 0 | 0 | 0.022 | 0.004 | 0.22 | 0 | 0 | 0 | 0 | 0.248 | 0 |
| CPM-N2 | 0.07 | 0 | 0 | 0.177 | 0.109 | 0.248 | 0 | 0 | 0 | 0 | 0.24 | 0.014 |
| CPM-N3 | 0.046 | 0 | 0 | 0.112 | 0 | 0.196 | 0 | 0 | 0 | 0 | 0.335 | 0 |
| CPM-P1 | 0.02 | 0 | 0.016 | 0 | 0.052 | 0.044 | 0 | 0 | 0 | 0 | 0.058 | 0 |
| CPM-P2 | 0.03 | 0.029 | 0.007 | 0.2 | 0 | 0.064 | 0 | 0.039 | 0 | 0 | 0.096 | 0.01 |
| CPM-P3 | 0.088 | 0 | 0.078 | 0.032 | 0.021 | 0.06 | 0 | 0 | 0 | 0 | 0.1 | 0 |

| Monocytes | Macrophages M0 | Macrophages M1 | Macrophages M2 | Dendritic cells resting | Dendritic cells activated | Mast cells resting | Mast cells activated | Eosinophils | Neutrophils | P-value | Pearson Correlation | RMSE |
|---|---|---|---|---|---|---|---|---|---|---|---|---|
| 0.198 | 0 | 0 | 0 | 0 | 0 | 0 | 0.039 | 0 | 0.225 | 0.010 | 0.417 | 0.922 |
| 0.056 | 0 | 0 | 0.026 | 0 | 0.017 | 0 | 0 | 0 | 0.04 | 0.010 | 0.524 | 0.856 |
| 0.268 | 0.002 | 0 | 0 | 0 | 0.006 | 0 | 0 | 0 | 0.036 | 0.010 | 0.556 | 0.832 |
| 0.753 | 0.025 | 0 | 0 | 0 | 0 | 0 | 0 | 0.003 | 0.029 | 0.010 | 0.643 | 0.766 |
| 0.464 | 0.037 | 0.002 | 0 | 0 | 0.002 | 0 | 0 | 0.02 | 0 | 0.010 | 0.650 | 0.770 |
| 0.574 | 0 | 0 | 0.029 | 0 | 0.008 | 0.008 | 0 | 0 | 0 | 0.010 | 0.704 | 0.724 |

**Supplemental Figure 2. Estimate of 22 immune cell types using CIBERSORT in PBMC samples from 3 uninfected and 3 COVID-19 participants.**

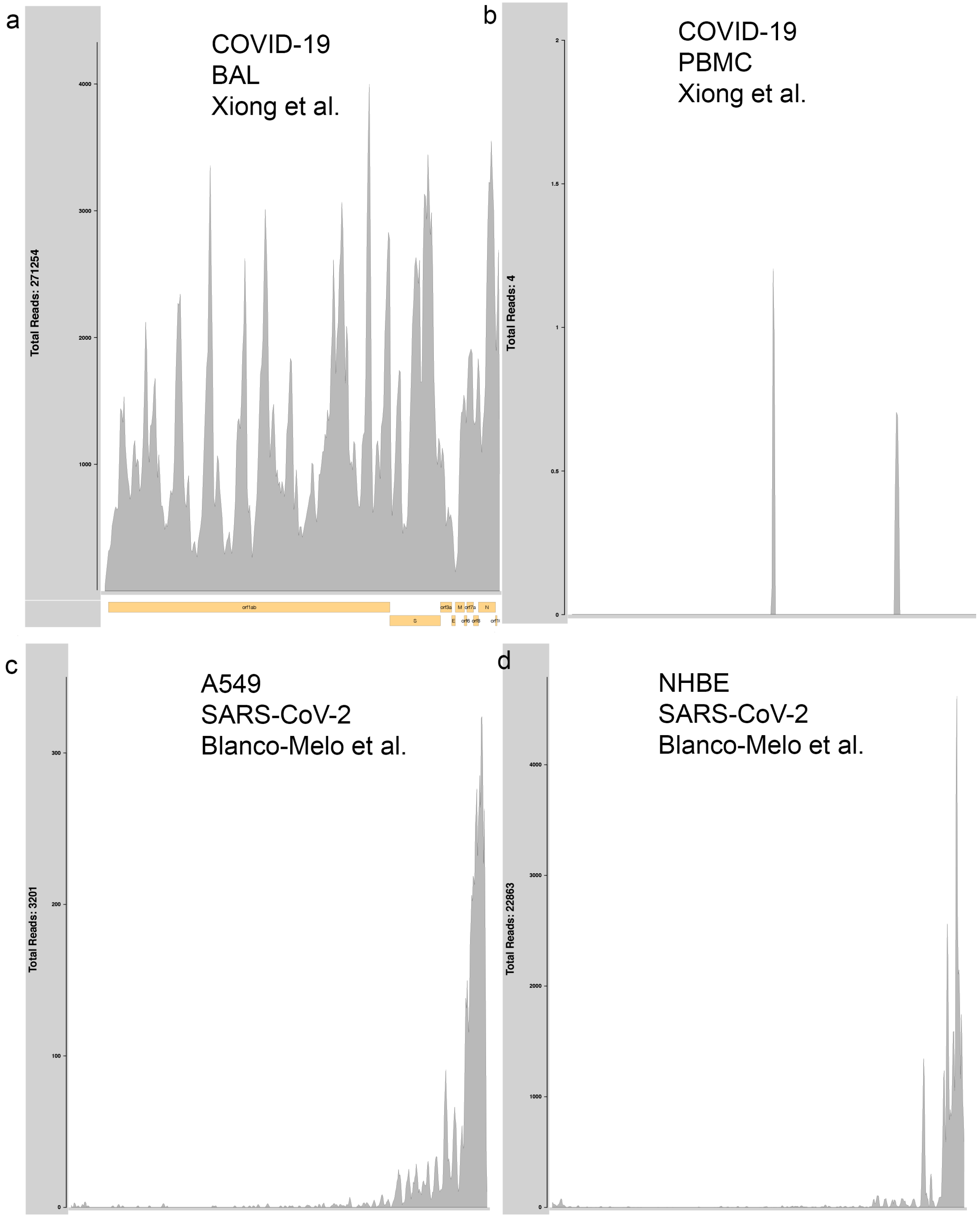


**Supplemental Figure 3. Mapping of RNA-Seq reads to SARS-CoV-2 reference in COVID-19 BAL, COVID-19 PBMC, A549 infected, and NHBE infected cells.**

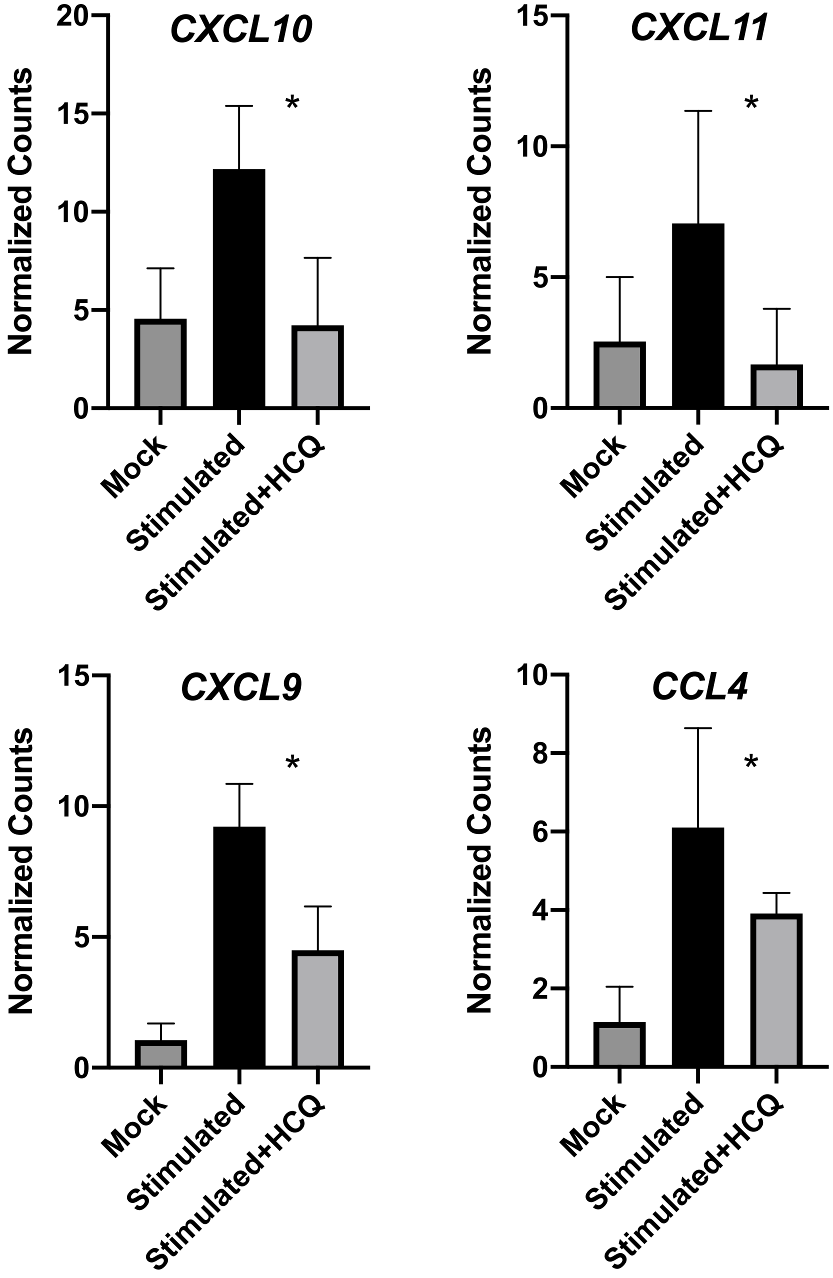


**Supplemental Figure 4. Normalized gene expression of chemokine genes in pDC mock treated, stimulated with RNA-IC, or stimulated and cotreated with HCQ.**

| Input Sample | B cells naive | B cells memory | Plasma cells | T cells CD8 | T cells CD4 naive | T cells CD4 memory resting | T cells CD4 memory activated | T cells follicular helper | T cells regulatory (Tregs) | T cells gamma delta | NK cells resting |
|---|---|---|---|---|---|---|---|---|---|---|---|
| CPM-Ctrl1 | 0.019 | 0 | 0 | 0.06 | 0 | 0.147 | 0 | 0 | 0 | 0 | 0.047 |
| CPM-Ctrl2 | 0.052 | 0 | 0 | 0 | 0 | 0.082 | 0 | 0.012 | 0.001 | 0 | 0.035 |
| CPM-Ctrl3 | 0.029 | 0 | 0 | 0.021 | 0 | 0.089 | 0.002 | 0 | 0.013 | 0 | 0.017 |
| CPM-nCoV1 | 0 | 0.023 | 0.062 | 0 | 0.064 | 0 | 0.017 | 0 | 0.031 | 0 | 0 |
| CPM-nCoV2 | 0 | 0.034 | 0 | 0.01 | 0.007 | 0 | 0 | 0 | 0.03 | 0 | 0.03 |

| NK cells activated | Monocytes | Macrophages M0 | Macrophages M1 | Macrophages M2 | Dendritic cells resting | Dendritic cells activated | Mast cells resting | Mast cells activated | Eosinophils | Neutrophils | P-value | Pearson Correlation | RMSE |
|---|---|---|---|---|---|---|---|---|---|---|---|---|---|
| 0 | 0.024 | 0.606 | 0.005 | 0.059 | 0 | 0 | 0.021 | 0 | 0 | 0.012 | 0.010 | 0.452 | 0.891 |
| 0 | 0 | 0.578 | 0.001 | 0.128 | 0.092 | 0 | 0.004 | 0 | 0 | 0.014 | 0.010 | 0.491 | 0.869 |
| 0 | 0.004 | 0.588 | 0.036 | 0.103 | 0 | 0 | 0.048 | 0 | 0.034 | 0.015 | 0.000 | 0.523 | 0.852 |
| 0.044 | 0.011 | 0.045 | 0.039 | 0.298 | 0 | 0.108 | 0.122 | 0 | 0.076 | 0.061 | 0.010 | 0.490 | 0.876 |
| 0 | 0 | 0.489 | 0.065 | 0.144 | 0 | 0 | 0.076 | 0 | 0.107 | 0.008 | 0.010 | 0.435 | 0.899 |

**Supplemental Figure 5. Estimate of 22 immune cell types using CIBERSORT in BAL samples from 3 uninfected and 2 COVID-19 participants.**

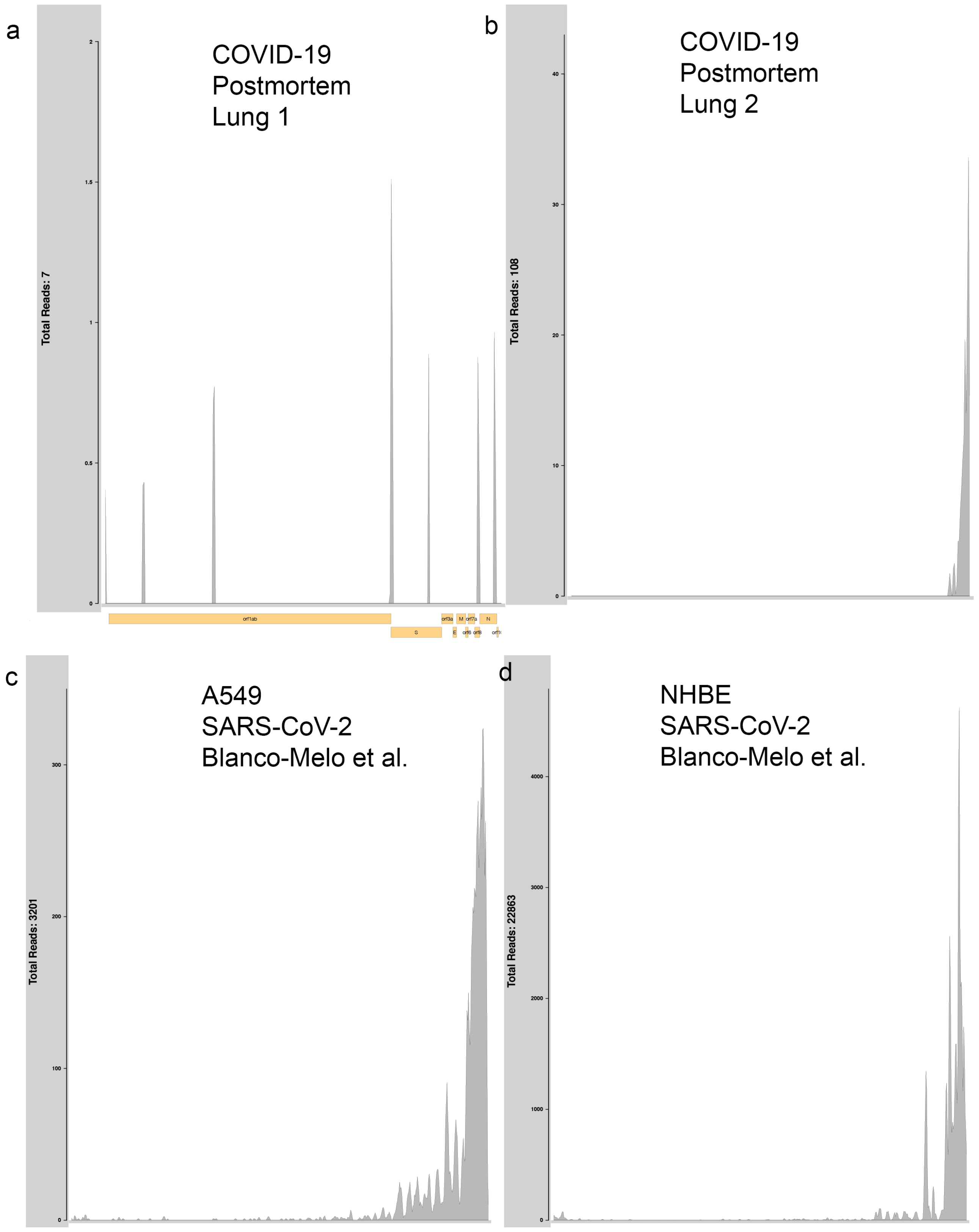


**Supplemental Figure 6. Mapping of RNA-Seq reads to SARS-CoV-2 reference in COVID-19 postmortem lung, A549 infected, and NHBE infected cells.**

| Input Sample | B cells naive | B cells memory | Plasma cells | T cells CD8 | T cells CD4 naive | T cells CD4 memory resting | T cells CD4 memory activated | T cells follicular helper | T cells regulatory (Tregs) | T cells gamma delta | NK cells resting | NK cells activated |
|---|---|---|---|---|---|---|---|---|---|---|---|---|
| Series15_HealthyLungBiopsy_2 | 0.051 | 0 | 0.014 | 0.088 | 0 | 0.143 | 0.012 | 0 | 0 | 0 | 0.08 | 0 |
| Series15_HealthyLungBiopsy_1 | 0.019 | 0 | 0.004 | 0.012 | 0 | 0.196 | 0 | 0 | 0 | 0 | 0.082 | 0 |
| Series15_COVID19Lung_2 | 0 | 0.002 | 0 | 0 | 0.009 | 0.03 | 0 | 0.006 | 0 | 0 | 0 | 0.042 |
| Series15_COVID19Lung_1 | 0 | 0.025 | 0 | 0 | 0.02 | 0.037 | 0 | 0.002 | 0.007 | 0.031 | 0 | 0.024 |

| Monocytes | Macrophages M0 | Macrophages M1 | Macrophages M2 | Dendritic cells resting | Dendritic cells activated | Mast cells resting | Mast cells activated | Eosinophils | Neutrophils | P-value | Pearson Correlation | RMSE |
|---|---|---|---|---|---|---|---|---|---|---|---|---|
| 0.094 | 0.08 | 0.013 | 0.209 | 0 | 0.03 | 0.06 | 0.004 | 0.039 | 0.083 | 0.010 | 0.211 | 0.997 |
| 0 | 0.13 | 0.065 | 0.011 | 0 | 0.109 | 0 | 0.371 | 0 | 0 | 0.010 | 0.279 | 1.013 |
| 0.063 | 0 | 0.044 | 0 | 0 | 0.126 | 0 | 0.45 | 0 | 0.228 | 0.000 | 0.596 | 0.814 |
| 0.066 | 0 | 0.043 | 0 | 0 | 0.03 | 0 | 0.587 | 0 | 0.129 | 0.000 | 0.597 | 0.829 |

**Supplemental Figure 7. Estimate of 22 immune cell types using CIBERSORT in postmortem lung samples from 2 uninfected and 2 COVID-19 participants.**

**Supplemental Table 1.**

| Dataset | Source | Notes |
|---|---|---|
| Hydroxychloroquine treated human PBMC cells | GSE74235 | Data from Untreated 24hr and HCQ 24 hr treated conditions was included in this analysis. |
| Hydroxychloroquine treated human primary macrophage cells | GSE95588 | Data from mono or co-culture TNF treated primary macrophages and mono or co-culture TNF hydroxycholoroquine treated primary macrophages was included in this analysis. |
| *In vitro* transcriptional response of SARS-CoV-2 infection | GSE147507 | Data from mock treated or SARS-CoV-2 infected A549 and NHBE cells was included in this analysis. |
| Transcriptome of patients infected with SARS-CoV-2 | PRJCA002326 | Data from bronchoalveolar lavage fluid (BALF) and peripheral blood mononuclear cells (PBMC) samples of COVID-19 patients was included in this analysis. |
| Hydroxychloroquine treated primary human plasmacytoid dendritic cells stimulated with RNA-IC | (Hjorton et al. 2018) | Data from four healthy individuals that had been either stimulated with RNA-IC for 6 hours only or stimulated with RNA-IC for 6 hours in the presence of HCQ was included in this analysis. |
| Transcriptome of postmortem lung biopsies of COVID-19 and uninfected individuals | GSE147507 | Data from two COVID-19 and two uninfected participants was included in this analysis. |

Hjorton, Karin, Niklas Hagberg, Elisabeth Israelsson, Lisa Jinton, Olof Berggren, Johanna K. Sandling, Kristofer Thörn, et al. 2018. "Cytokine Production by Activated Plasmacytoid Dendritic Cells and Natural Killer Cells Is Suppressed by an IRAK4 Inhibitor." *Arthritis Research & Therapy* 20 (1): 238.
